# Metabolic Nutrient Preferences of *Vibrio mimicus*: Leveraging Nucleotides and Oligopeptides from Yellow Catfish for Enhanced Infectivity

**DOI:** 10.1101/2023.07.27.550872

**Authors:** Yang Feng, Jiao Wang, Wei Fan, Yi Geng, Xiaoli Huang, Ping Ouyang, Defang Chen, Hongrui Guo, Huidan Deng, Weimin Lai, Zhicai Zuo, Zhijun Zhong

## Abstract

In the context of host-microbe interactions, the microenvironment plays a critical role in facilitating microbial survival, and variations in these microenvironments may influence the pathogenicity of microorganisms. *Vibrio mimicus*, a major pathogen responsible for infections in aquatic animals, poses a substantial threat to yellow catfish (*Pelteobagrus fulvidraco*) and grass carp (*Ctenopharyngodon idella*), two naturally occurring hosts displaying markedly different susceptibility levels. This study aims to unravel the underlying mechanisms behind this susceptibility discrepancy in the two teleost species. Employing metabolomic analysis, we identified a distinctive microenvironment in yellow catfish, characterized by abundant purine nucleotides and oligopeptides. Furthermore, a total of 67 specific metabolites were identified from both yellow catfish and grass carp, with 33 displaying heightened expression on the body surface of yellow catfish, including nucleotides, amino acids, and gangliosides, while 34 were predominantly expressed on the body surface of grass carp, primarily comprising lipids. Subsequent investigations revealed that certain compounds related to nucleotides and oligopeptides exhibited significant growth-promoting effects and were utilized by *V. mimicus* as nutrients, with deoxyguanosine proving to be notably more than twice as effective as glucose. Moreover, during *V. mimicus* infection, numerous metabolites such as oligopeptides, purine nucleotides, and specific metabolites experienced considerable depletion in the skin of yellow catfish. Concurrently, several genes associated with nucleosidase and peptidase were upregulated in the skin and muscles of infected fish. These findings suggest that the microenvironment provided by different hosts plays a pivotal role in determining the infectivity of the pathogen. Additionally, our results indicate that the microenvironment on the surface of yellow catfish, characterized by an abundance of purine nucleotides and oligopeptides, indirectly enhances *V. mimicus* growth, ultimately augmenting its infectivity.

## 1 Introduction

Bacterial infection is a multifaceted phenomenon that involves various factors such as virulence, host immunity, and bacterial living microenvironments. The significance of the microenvironment in which bacteria grow, interacting directly with their surroundings, cannot be underestimated, as it plays a crucial role in their survival and pathogenicity (Wessel et al., 2013). While studies often consider the impact of host immunity and virulence on bacterial infections, the microenvironment aspect is frequently overlooked. In the human body, a staggering number of symbiotic bacteria, approximately 3.8×10^13^, coexist with the host in a delicate balance (Sender et al., 2016). These symbiotic bacteria receive a rich supply of nutrients, which also benefits pathogenic bacteria. The composition of the host’s metabolites exerts complex effects on bacterial growth rates, as suggested in studies by Bonnet et al. (Bonnet et al., 2020). Consequently, a conducive microenvironment can potentially exacerbate bacterial infectivity, contributing to disease progression. To thrive and reproduce, microbial communities have evolved mechanisms to resist foreign threats and acquire nutrients through virulence factors and tools like hemolysin and protease. The balance of these factors in a suitable microenvironment determines the outcome of bacterial culture, which, if disturbed, can lead to excessive bacterial growth, accumulation of virulence factors, and, ultimately, disease and death in the host.

Despite the importance of the host nutritional microenvironment in bacterial pathogenicity, there has been a lack of extensive research in this area. Thus, in this study, we aim to explore the potential role of the host microenvironment in enhancing pathogen infectivity. To achieve this, we employ the yellow catfish as a model organism infected with *Vibrio mimicus*, a pathogenic species within the *Vibrio* genus. *Vibrio mimicus* shares morphological and physiological characteristics with *Vibrio cholerae*, causing symptoms similar to enteritis and sepsis in humans, much like *V. cholerae* non-O1 and O139 strains (Thompson et al., 2008). In freshwater environments, *V. mimicus* exhibits high pathogenicity in fish and crustaceans (Gao et al., 2015; Y. Geng et al., 2014; Jin-yu et al., 2000; Mustafa, 2016), particularly in Siluriformes, which include southern catfish, yellow catfish, and channel catfish (Fu et al., 2021; Y Geng et al., 2014). These fish species are highly susceptible to *V. mimicus* infection, leading to mortality rates exceeding 90% within 5-7 days of infection (Y Geng et al., 2014).

*Vibrio mimicus* has been found to possess numerous virulence factors akin to those of *V. cholerae*, suggesting a considerable degree of pathogenicity (Yu et al., 2020). Nevertheless, the underlying factors contributing to the heightened pathogenicity of *V. mimicus* in Siluriformes remain elusive. Siluriformes are well-regarded for their comparatively robust resistance to diseases commonly prevalent in sewage environments. Lau et al. (Lao, 2017) reported an LC_50_ of 3.17 × 103 CFU/mL when yellow catfish were exposed to *V. mimicus*. Conversely, Li and Fu (Fu, 2020; Y.-W. Li et al., 2019) reported relatively higher LC_50_ values of 3.42 × 10^5^ CFU/mL and 1.3 × 10^6^ CFU/mL for channel catfish and yellow catfish, respectively. Additionally, a separate study found that the LC_50_ of *V. mimicus* for grass carp, another bony fish, was 1.076 × 10^7^ CFU/mL (Zhang, 2013), significantly surpassing that of Siluriformes. Interestingly, crustaceans, lacking adaptive immune systems (Huang & Ren, 2020; Vazquez et al., 2009), also demonstrated heightened tolerance to *V. mimicus*, with an LC_50_ of 2.09 × 10^7^ CFU/mL observed for *Penaeus monodon* (Raja et al., 2017) and 3.28 × 10^5^ CFU/mL for *Macrobrachium nipponense* (Jiang et al., 2022). It proves intriguing to elucidate the factors underpinning the heightened pathogenicity of *V. mimicus* in Siluriformes, particularly with a directed emphasis on the yellow catfish. Through an in-depth examination of the metabolite composition within the yellow catfish, a prominent fishery product in China (Bureau of Fisheries, 2022), our objective is to unveil novel perspectives on how the host microenvironment may augment *V. mimicus* infectivity. This endeavor additionally yields invaluable insights into bacterial pathogenesis and potentially paves the way for therapeutic interventions.

## 2 Materials and Methods

### 2.1 Bacteria

The strain *V. mimicus* SCCF01 was isolated from diseased yellow catfish at a commercial aquaculture site in Sichuan province, China, in 2014 (Y Geng et al., 2014). Employing aseptic techniques, the strain was isolated from the infected fish and subsequently inoculated onto Luria Broth (LB), then incubated at 28 ℃ for 24-48 hours. Following this, single colonies with uniform morphology and size were meticulously chosen and cultured to obtain pure strains, which were later identified through genome analysis (Yu et al., 2020). For metabolite analysis and animal challenge trials, *V. mimicus* strains were cultured in a BHI medium at 28 ℃ for 24 hours.

### 2.2 Establishment of Median Lethal Concentration (LC_50_) Model

The experimental setup for establishing the LC_50_ model involved the utilization of specific animal species, namely, the mouse (*Mus musculus*) KM-SPF and aquatic organisms, grass carp (8-9 cm) and red crayfish (*Procambarus clarkia*). The mice, with an average weight of 35.5 ± 3.5 g, were procured from Chengdu Dossy Experimental Animals Co., Ltd, whereas the grass carp and red crayfish, also of similar weight, were obtained from a commercial farm located in Wenjiang, Sichuan, China.

To ensure the well-being of the animals during the experiment, they were acclimated for a period of two weeks and provided with commercial pellets twice daily. For the mice, the ambient air temperature was regulated at 22 ± 2°C, and they were supplied with mineral water and paulownia shavings as bedding, while the aquatic animals were maintained at a water temperature of 22 ± 2°C. Furthermore, the aquatic environment was carefully monitored, with ammoniacal nitrogen and nitrite levels maintained at 0–0.02 mg/L and pH levels at 7.0–7.5. Ethical considerations were meticulously adhered to throughout the study, in accordance with the guidelines for animal experiments stipulated by Sichuan Agricultural University. The study received approval from the university’s Animal Care and Use Committee, under permit number 2020103001.

The experiment comprised six groups of animals: a control group administered with 0.65% and 0.9% physiological saline for aquatic animals and mice, respectively, and five experimental groups exposed to varying concentrations of purified *V. mimicus*, ranging from 10^5^ to 10^9^ CFU/mL for aquatic animals and 10^6^ to 10^10^ CFU/mL for mice. Each animal in the experimental groups underwent a specific challenge procedure, involving a 0.1 mL intramuscular injection in mice or immersion in the *V. mimicus* solution for 30 minutes in the case of aquatic animals. Subsequently, gross lesions and mortalities were closely monitored and recorded on an hourly basis, with immediate removal of deceased animals. The LC50 of *V. mimicus* for the animals was determined using the Probit method, yielding essential insights into the pathogenicity of the strain and its impact on the respective animal species.

### 2.3 Samples for Metabolites Comparison

In this study, the analysis of metabolites involved the classification of samples into five distinct groups, each consisting of four replicates. Bacterial samples, specifically *V. mimicus*, were subjected to metabolite analysis after being incubated in four different media, namely LB, BHI, TSB, and DMEM, at 28°C for 24 hours to minimize the potential impact of the medium on the bacteria. Subsequently, the resulting assemblages were washed three times with cold PBS and then stored at –80℃ under nitrogen for subsequent analysis. Yellow catfish (weighing 69.6±6.8 g) and Grass carp (weighing 78.3±5.3 g) were procured from a commercial farm located in Wenjiang. These fish were acclimated for a period of two weeks, and their skin and muscle tissues were collected from below the anterior side of the dorsal fin and above the lateral line. For further analysis, the tissue of every three fish was homogenized into a single sample and then stored at –80℃ under liquid nitrogen. By following this meticulous procedure, we ensured a robust and reliable comparison of metabolites among the different groups, as well as accurate data to support our research findings.

### 2.4 *V. mimicus* Infected Model

The *V. mimicus* Infected Model was utilized in this study to investigate the effects of bacterial infection on healthy yellow catfish. Yellow catfish, which exhibited complete appendages, showed no disability, and possessed robust vitality, were chosen randomly and divided into two groups: the control group and the challenge group. The control group received treatment with 0.65% physiological saline, while the challenge group was treated with 1.0 × 105 CFU/mL purified strains of *V. mimicus*. Prior to the challenge, the fish underwent a 30-minute immersion in pre-trialed water before being transferred to fresh water. Throughout the experiment, the water temperature was maintained at 23 ± 2°C, and the pH level was carefully regulated within the range of 7.5–8.5. The yellow catfish in both groups were fed with commercial feed, equivalent to 5% of their body weight, daily between 6:00 and 7:00 pm. The observation period lasted for seven days, during which the researchers recorded the mortality rate and activity level of the fish in each group. Seven days post-challenge, four skins from the yellow catfish in each group were randomly collected for subsequent analysis of metabolites and RNA-seq.

### 2.5 Metabolite Analysis using LC-MS Detection

Metabolic profiling was conducted on the samples following the protocol as described below: Each sample was mixed with an internal standard in a 1.5 mL Eppendorf tube, followed by the addition of an ice-cold mixture of methanol and water (4:1 v/v). Subsequently, the samples were stored at –20°C for 5 minutes, ground at 60 Hz for 2 minutes, ultrasonicated in an ice water bath for 10 minutes, and then held at –20°C for 30 minutes. The extract was then centrifuged at 13000 rpm and 4°C for 15 minutes. The resulting supernatant (300 μL) was dried using a freeze-concentration centrifugal dryer and subsequently reconstituted with a 400 μL mixture of methanol and water (1:4, v/v) with 30 seconds of vortexing. The reconstituted samples were kept at –20°C for two hours, followed by centrifugation at 13000 rpm and 4°C for 10 minutes. The resulting supernatants (150 μL) were collected using crystal syringes, filtered through 0.22 μm microfilters, and transferred to LC vials. The vials were stored at –80°C until LC-MS analysis. To create pooled quality control (QC) samples, aliquots from all 18 samples were mixed. LC-MS analysis was conducted using a Dionex Ultimate 3000 RS UHPLC system coupled with a Q Exactive quadrupole Orbitrap mass spectrometer equipped with a heated electrospray ionization (ESI) source (Thermo Fisher Scientific, Waltham, MA, USA) in both ESI positive and negative ion modes. The ACQUITY UPLC HSS T3 column (100 mm × 2.1mm, 1.8 μm) was used for both positive and negative methods. QC samples were regularly injected throughout the analytical run to assess repeatability.

Metabolites were identified using progenesis QI Data Processing Software (Waters Corporation, Milford, USA) and public databases like the Human Metabolome Database (http://www.hmdb.ca/). Correlation analysis was performed using the Pearson correlation coefficient to measure the linear correlation between two quantitative variables, and the P value was utilized to indicate correlation significance.

Given that the samples of *V. mimicus*, grass carp, and yellow catfish were independently selected from different populations during the early stage of the experiment, the independent samples Kruskal-Wallis rank-sum test was employed to examine the differences in metabolite composition among groups (*P*<0.05) for screening differential metabolites. Meanwhile, the unpaired Student’s t-test was applied to detect the differences in data for the metabolomic analysis of yellow catfish skin after *V. mimicus* infection. Differential metabolites were selected based on a combination of statistically significant thresholds of variable influence on projection (VIP) values obtained from the (orthogonal) partial least-squares-discriminant analysis (OPLS-DA) model, where metabolites with VIP values greater than 1.0 and P values less than 0.05 were considered differential metabolites. Subsequently, differential metabolites were annotated with metabolic pathways using the Python software package scipy.stats (KEGG database: www.kegg.jp/kegg/pathway.html), and the pathways involving the differential metabolites were identified. All pathways underwent enrichment analysis, and the biological pathways most relevant to the experimental treatment were identified through Fisher’s exact test.

### 2.6 Effect of Compounds on *V. mimicus*

The compounds utilized in this study were acquired from a local company, and Table 1 displays their respective purity levels. The SCCF01 strain was cultivated in LB liquid medium at 30°C for eight hours, and the resulting bacterial suspension was diluted with fresh LB liquid medium to achieve an OD_600_=1.0 for subsequent experimentation. To assess the impact of these compounds on bacterial growth, varying concentrations were incorporated into the LB medium (at a 50% concentration). The SCCF01 strain, with an initial OD_600_ of 1, was then introduced at 1% (v/v) into each liquid medium. Subsequently, the cultures were incubated at 30°C and 180 r/min, and samples were collected at 0h and 24h to measure OD_600_ values.

**Table 1.**
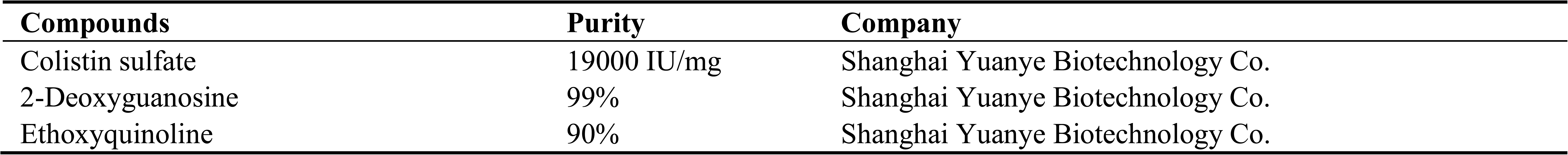
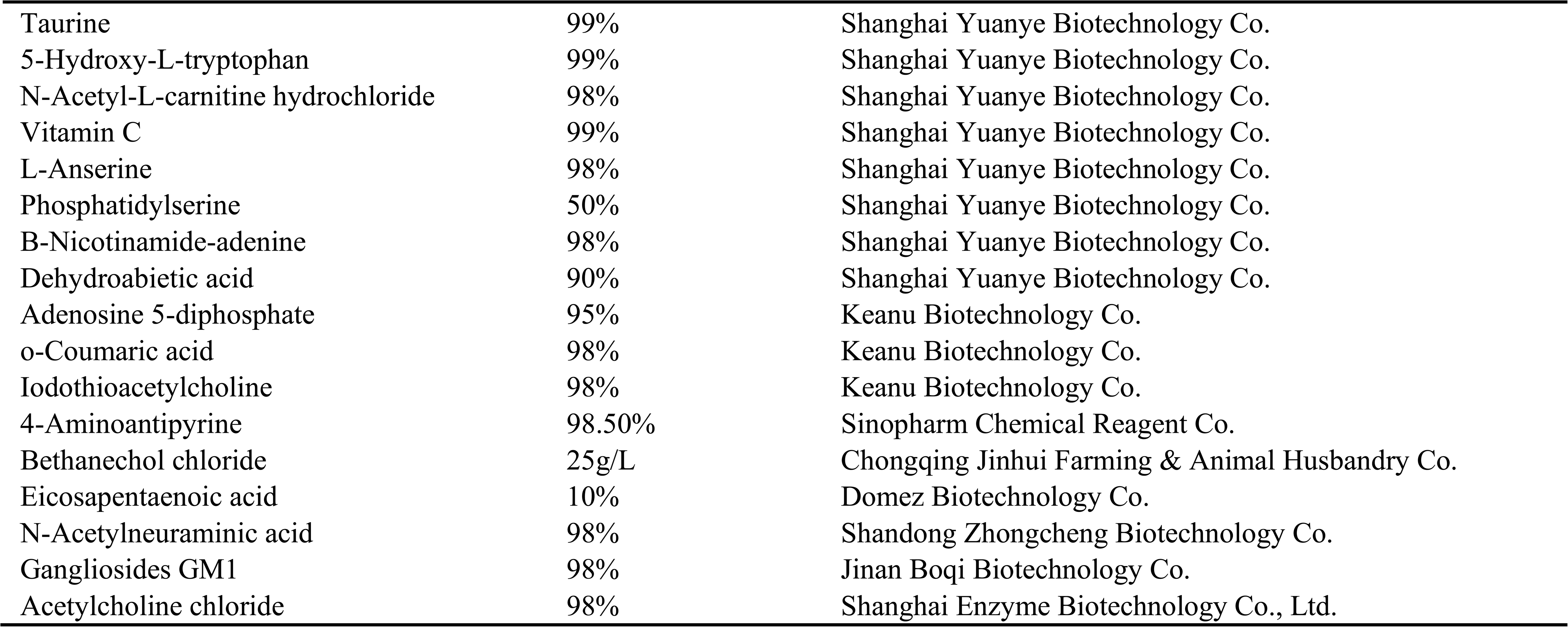
Information about compounds.

To evaluate carbon source utilization capacity, different compounds (at concentrations of 1.0 mg/L and 2.0 mg/L) were added as carbon sources to M9 basal medium containing five mM CaCl_2_ and 32 mM MgSO_4_. As a control group, glucose was added at the same concentrations (1.0 mg/L and 2.0 mg/L). Again, the SCCF01 strain, with an initial OD_600_ of 1, was inoculated at 1% (v/v) into each liquid medium, and the cultures were incubated at 30°C and 180 r/min. Samples were collected at 0h and 24h to measure OD_600_ values.

For investigating motility, LB agar plates with 0.5% agar content (w/v) were prepared, and various compounds were added to maintain a concentration of 500 mg/L in the plates. A bacterial suspension (20 μL of SCCF01 with an initial OD_600_ of 1.0) was then vertically inoculated into blank susceptibility Test Discs, each with a diameter of 6 mm. Each compound’s test was repeated four times. The inoculum was allowed to dry naturally, and the LB agar plates were incubated at 28°C for 24h. Subsequently, the diameter of the bacterial colonies was measured using vernier calipers.

### 2.7 RNA-seq Analysis

In this study, we conducted RNA sequencing on four skin and muscle samples in each experimental group. The sample preparation and data analysis procedures were previously described in a study conducted by Feng et al. (Feng et al., 2023). The resulting data was submitted to the NCBI and is publicly accessible under the accession number PRJNA938908. To analyze the unigenes, we employed the BLAST software to compare them to pathways, such as “Nucleotide metabolism” and “Amino acid metabolism.” The expression levels of differentially expressed genes (DEGs) were determined using Transcripts Per Million (TPM) values, and DESeq2 algorithms were utilized for the selection of DEGs. The P values were corrected using the Benjamini-Hochberg method, with a screening threshold of FDR ≤ 0.05 and an absolute value of log2 Fold Change ≥ 2.

To validate the transcriptome, we performed qPCR using a TaKaRa kit and a BioRad Thermo Cycler. The primer sequences for each gene used in this analysis can be found in Table 2. The reactions were carried out in a ten μL mixture containing SYBR Green PCR Master Mix, diethylpyrocarbonate-treated water, forward and reverse primers, and cDNA. The samples underwent 40 cycles of amplification, and we employed the 2^−ΔΔCT^ method to calculate the relative changes in mRNA expression based on the qPCR results (Feng et al., 2023).

**Table 2.**
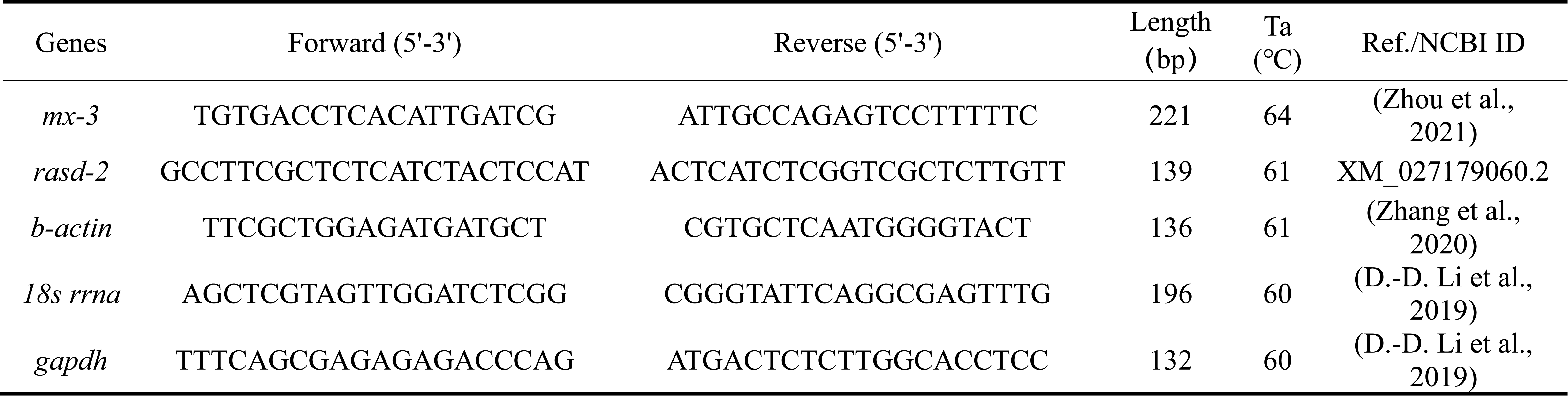
Primers of various genes detected with qPCR.

### 2.8 Statistical Analysis

The mean value and standard deviation were utilized to express the obtained results. The Student’s t-test was conducted to ascertain the statistical significance of intergroup disparities (SPSS v.20.0, IBM Corp., Armonk, New York, USA). Significance was attributed to results with a *P*-value less than 0.05, while results with a *P*-value less than 0.01 were considered highly significant.

## 3 Results

### 3.1 Susceptivity of Animals to *V. mimicus*

In this study, a series of infected animals with *V. mimicus* were carefully selected, with mice serving as mammalian surrogate models for human infection. The primary objective was to assess the LC_50_ analysis of SCCF01 on these animals. The results unveiled that *V. mimicus* displayed varying degrees of susceptibility among different animal species, exhibiting high toxicity to Siluriformes, followed by *P. clarkii* and grass carp. However, its toxicity to mice was comparatively weak (Fig. 1 A). Upon evaluating the clinical manifestations, it was observed that *V. mimicus* induced skin and muscle ulceration symptoms in both Siluriformes and grass carp (Fig. 1 B-C). Additionally, in yellow catfish, there were indications of necrotizing myositis, epidermal necrosis, and serous epidermal inflammation, with numerous *V. mimicus* parasites found in the dermal tissue (Fig. 1 D-G). Remarkably, other organs, such as the spleen, liver, and kidney, exhibited relatively minor damage, and typical symptoms of gastroenteritis, often associated with *V. mimicus* infections, were rarely observed in yellow catfish. Based on these noteworthy findings, we hypothesize that yellow catfish possess unique substances, distinct from those present in grass carp (Fig. 1H). These substances seem to facilitate the parasitism and reproduction of *V. mimicus* within the skin of yellow catfish.

**Fig. 1.**
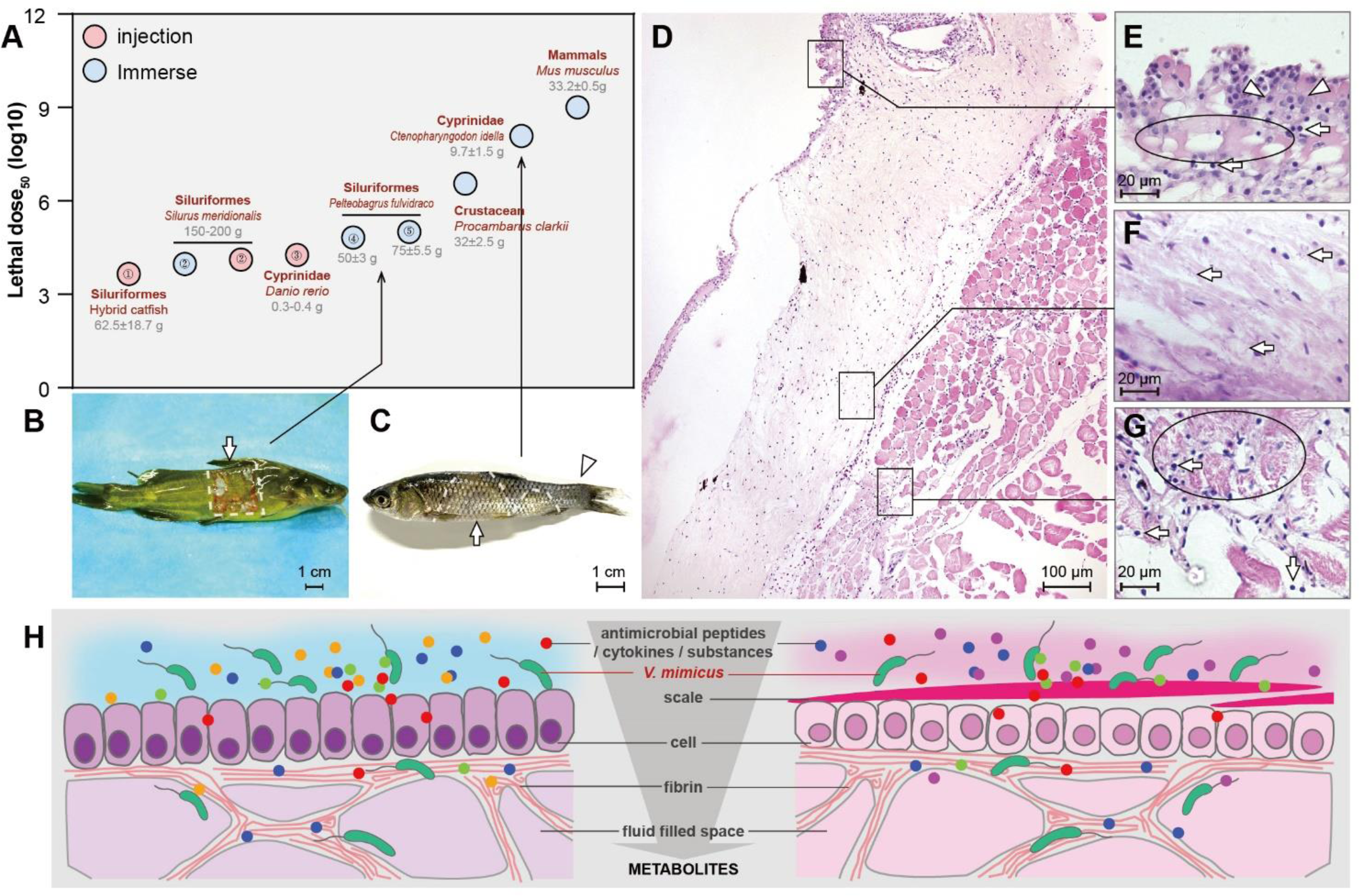
Depicts the general characteristics of animals under *V. mimicus* infection. A: The LC_50_ of *V. mimicus* to different animals. B: the general lesion of *V. mimicus* infection in yellow catfish, with a square ulcer indicated by an arrow. C: the gross lesion of *V. mimicus* infection in grass carp, with arrows indicating discolored and ulcerated skin and an arrowhead indicating a rotten tail. D: severe dermatomyositis in yellow catfish infected with *V. mimicus* as observed in histopathology. E, F, and G: the enlargement of the epidermal layer, dermis, and muscle of yellow catfish, respectively. Neutrophil infiltration, serous cell hyperplasia, cloudiness bacteria formed by *V. mimicus*, and muscle necrosis are indicated by arrows, circles, and arrowheads as appropriate. H: the pattern of the surface microenvironment of yellow catfish and grass carp.

### 3.2 Metabolic Composition of Yellow Catfish and Grass Carp

In this study, we delved into the microenvironment of the yellow catfish body surface, investigating its significance as the preferred habitat for *V. mimicus*. To achieve this, we conducted a comparative analysis of the metabolite composition in the skin and muscle of yellow catfish and grass carp (Fig. 2A). The samples were collected from the dorsal fin base. The mass spectral signals were collected and preprocessed using both positive and negative ion scanning modes (Fig. 2B), leading to the identification of 7679 (pos+) and 5685 (neg-) effective peaks, respectively (Fig. 2C). Among these peaks, 1213 metabolites were successfully identified and annotated through the KEGG pathway, KEGG compounds, and HMDB compounds database (Fig. S1). The PCA analysis revealed excellent intra-group correlation and noticeable inter-group differences across the various samples (Fig. 2E).

**Fig. 2.**
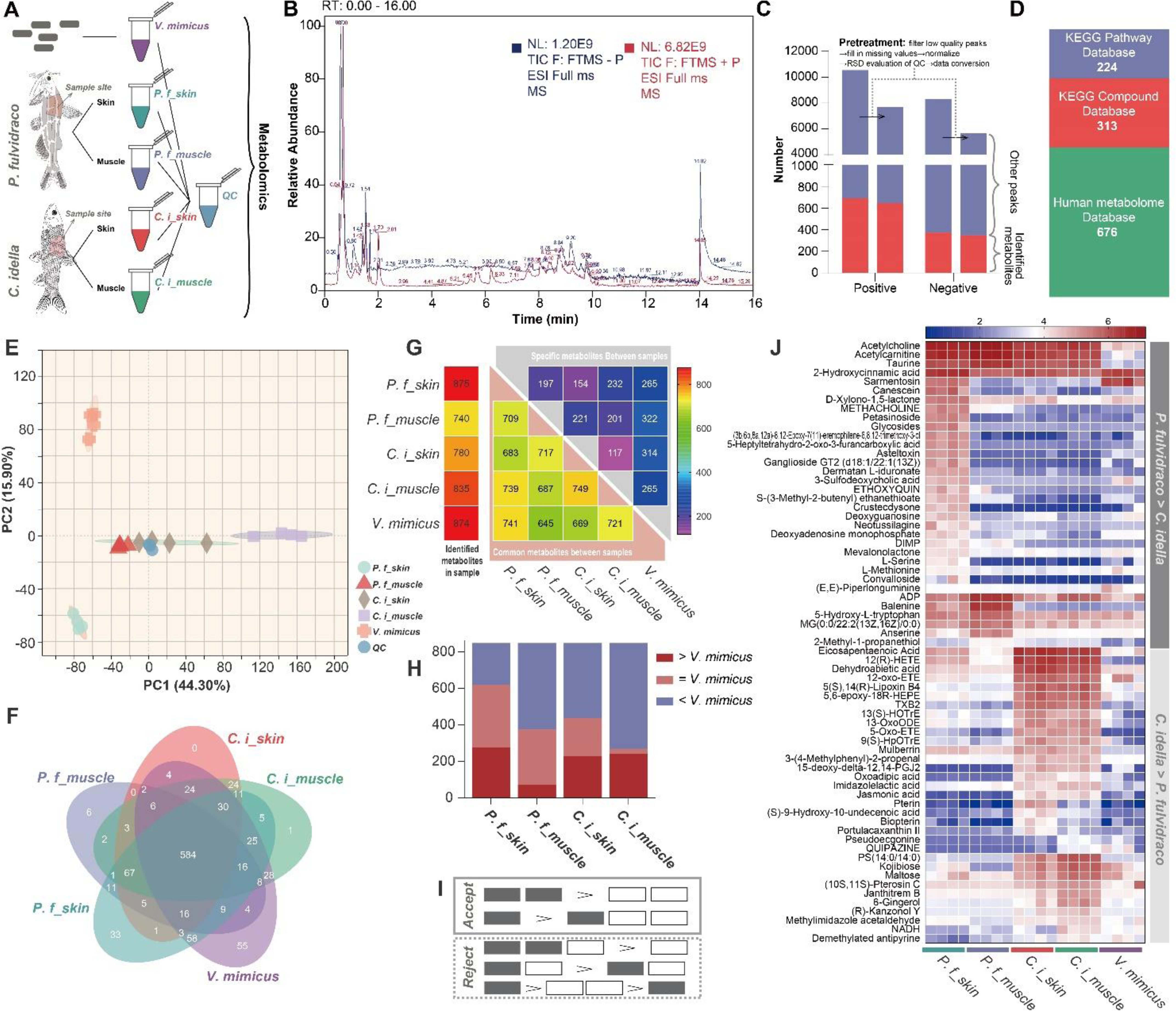
The metabolomic analysis and metabolite screening of different samples. A: the types and collection criteria of samples used for LC-MS. B: the ESI (+, –) ion flow diagram of quality control samples. C: the identification and statistics of total ions. D: the annotation information of total ions in three databases. E: the PCA analysis of different samples. F: the Venn diagram of annotated metabolites between samples. G: the comparison of identified metabolites in different samples. H: the comparison of co-annotated metabolite abundance in all samples. I: the standard for screening target metabolites in two fish species, where the abundance of metabolites was statistically analyzed and expressed as “>,” “<,” or “=.” J: the expression abundance of 67 candidate metabolites in different samples.

Through our comparison, we found 584 annotated metabolites from different samples (Fig. 2F). Interestingly, the skin of yellow catfish shared the most common annotations with *V. mimicus*, followed by grass carp muscle, as determined through statistical analysis (Fig. 2G). Notably, the total amount of metabolites equal to or significantly higher than those found in *V. mimicus* was much greater in the skin of yellow catfish compared to other samples (Fig. 2H). This observation suggests that the metabolites in the skin microenvironment of yellow catfish potentially provide a more suitable substrate for *V. mimicus* survival. Furthermore, when comparing the four types of fish tissues, *V. mimicus* exhibited higher levels of short peptides and nucleotide substances, along with lower levels of lipid metabolites. Interestingly, the surface composition of yellow catfish and grass carp corresponded to one of these features.

In our investigation, we also identified specifically high-expressed metabolites in yellow catfish and grass carp (Fig. 2I). To put it simply, the strategy depicted in Fig. 2I allowed us to pinpoint metabolites that exhibited significant upregulation within each species. Through this protocol, a total of 67 specifically expressed metabolites were found in both fish species (Fig. 2J, Table 3), indicating a clear interspecific correlation (Fig. S2). The specifically expressed metabolites in yellow catfish included nucleotides, amino acids, and other metabolites, whereas those in grass carp mainly comprised fatty acids and other metabolites (Table 3). Additionally, our study unveiled 172 metabolites with unknown functions, which offered valuable insights into the oligopeptides highly expressed in the skin of yellow catfish (Fig. 3).

**Fig. 3.**
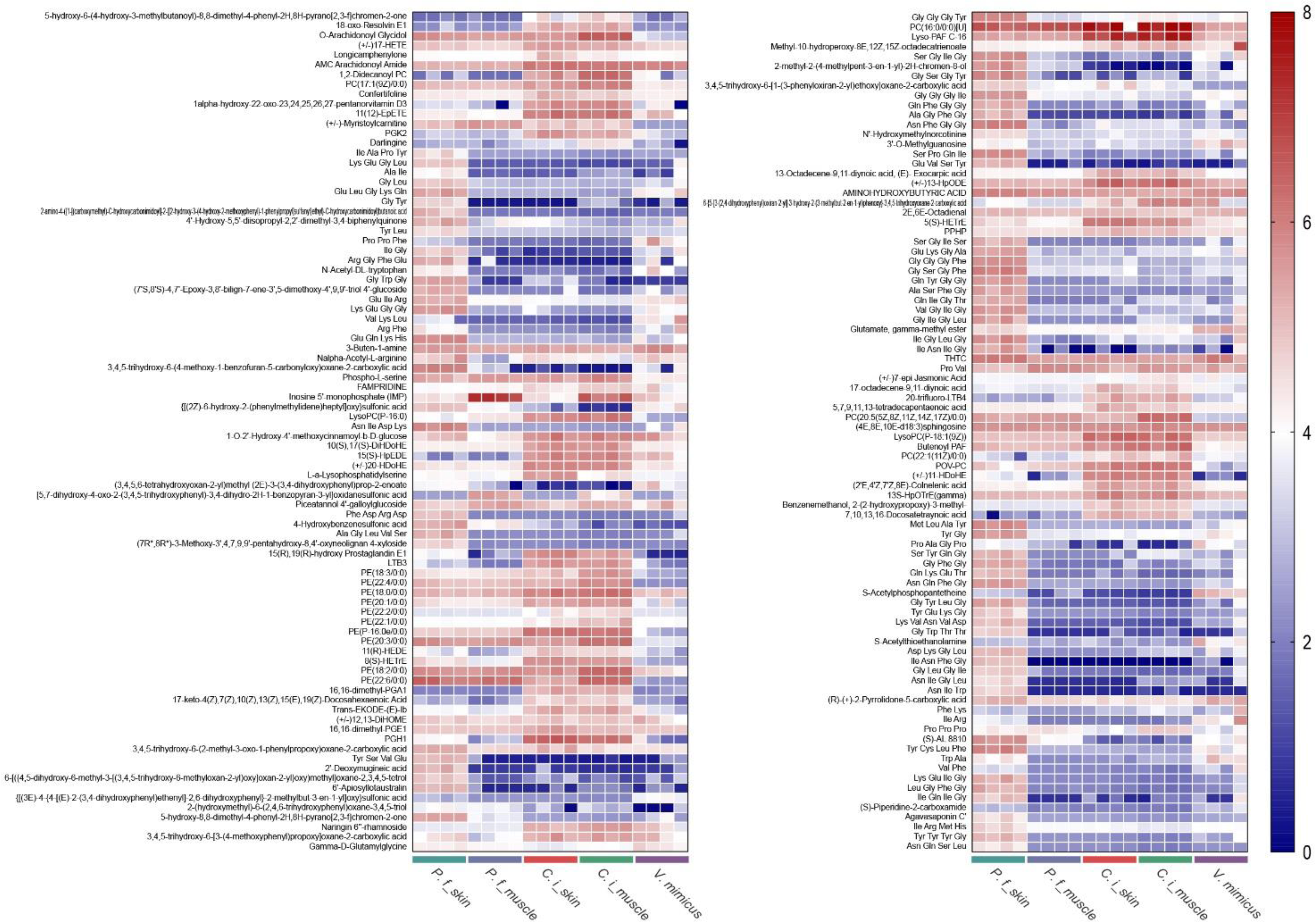
The expression abundance of unknown functional metabolites in different samples.

**Table 3.**
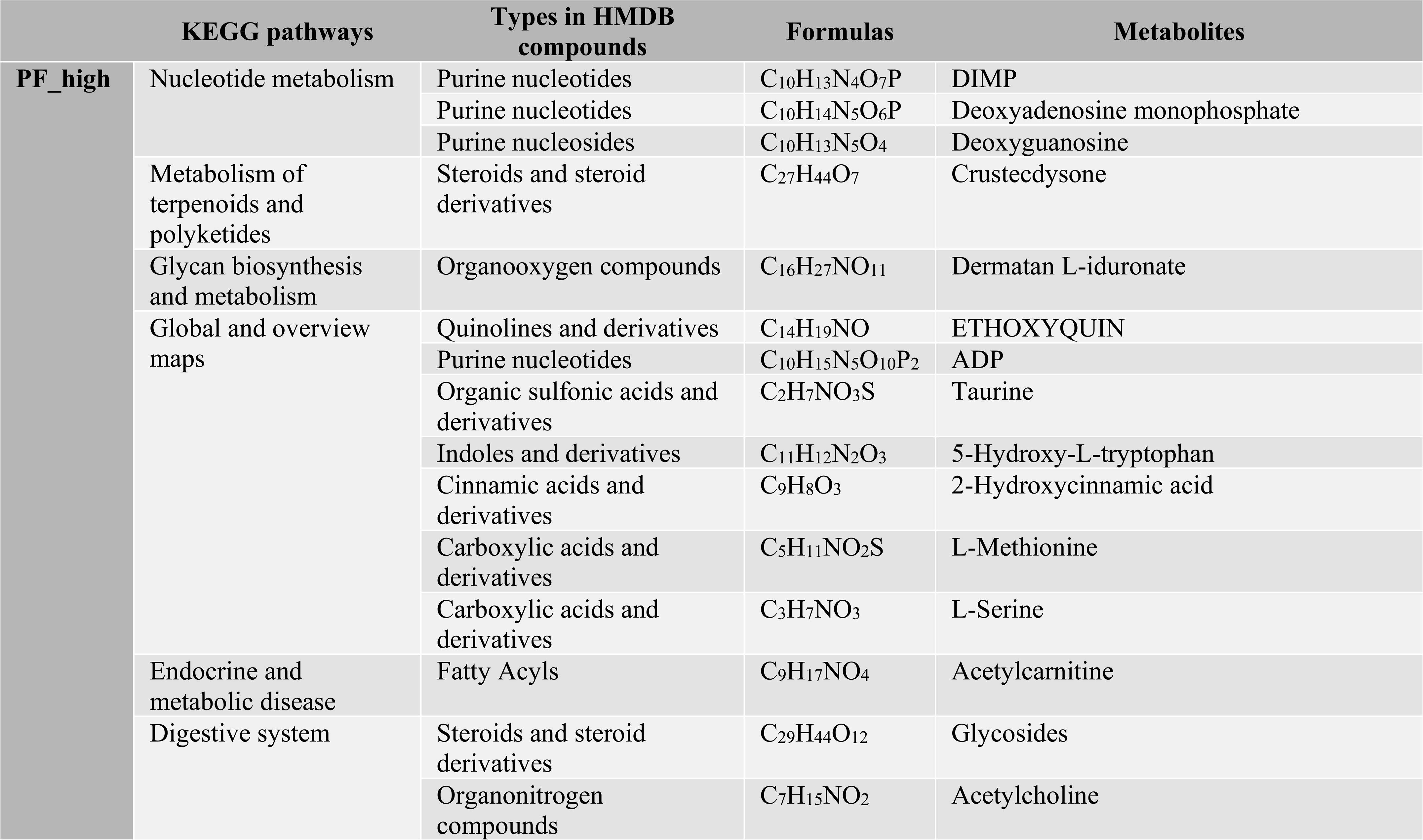
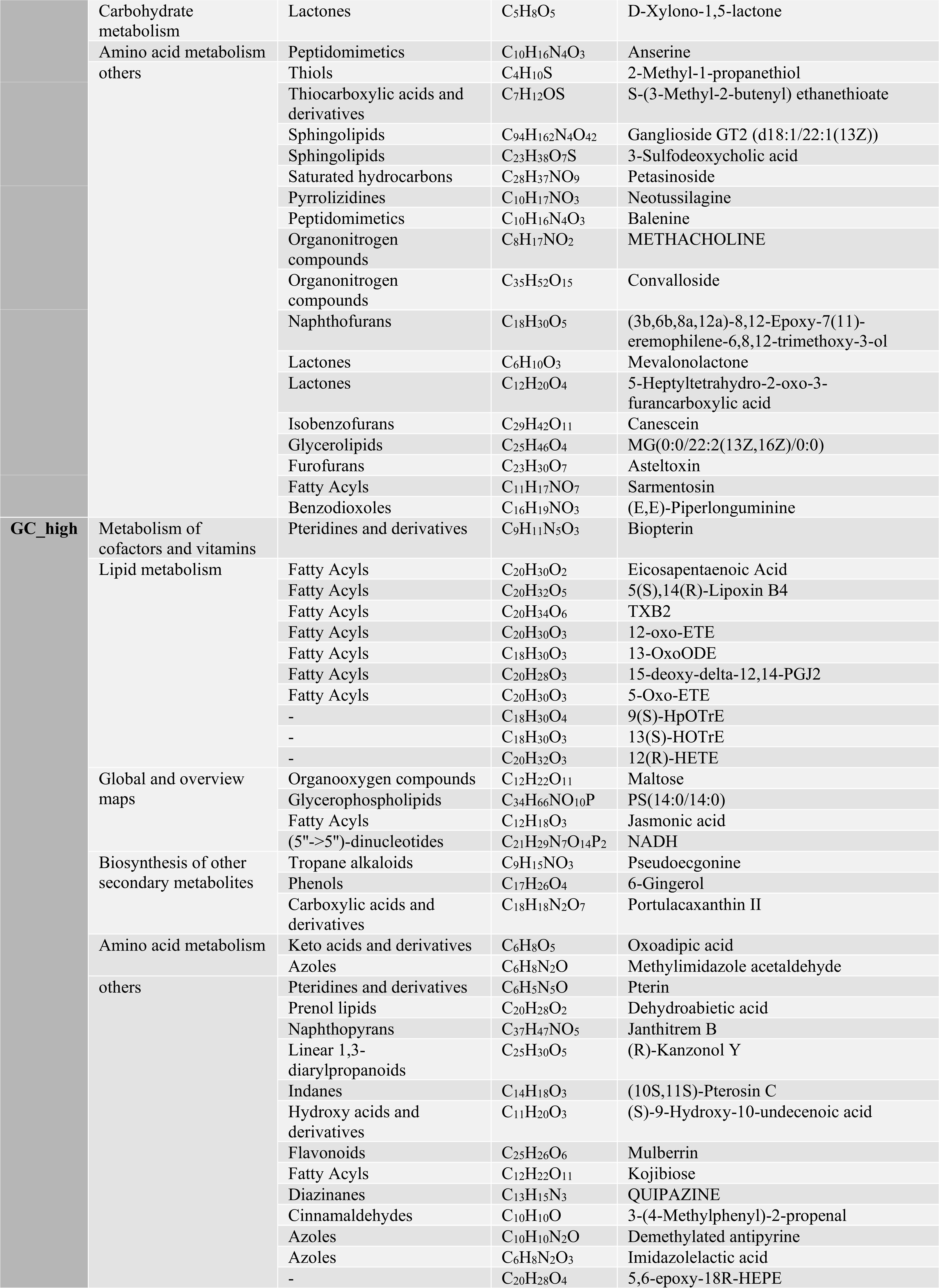
Relevant information of 67 target metabolites screened in this study.

### 3.3 Life Effects of Candidate Metabolites on *V. mimicus*

Besides, we investigated the effects of specific compounds or analogs on both the growth and carbon source utilization of *V. mimicus*. To achieve this, we carefully selected 20 compounds or analogs from a pool of 67 metabolic categories, comprising 15 highly expressed metabolites in yellow catfish and five in Grass carp (Fig. 4 A, Fig. S3). We then assessed the impact of these compounds on the growth of *V. mimicus* at different concentrations. Interestingly, some of the compounds displayed a biphasic effect on the growth of *V. mimicus*. At low concentrations, these compounds promoted growth, while at higher concentrations, they inhibited growth. Notable examples include N-Acetylneuraminic acid, Ganglioside GM1, vitamin C, (R)-3-(Acetyloxy)-4-(trimet-hylammonio) butyrate hydrochloride (Acetyl L-Carnitine), β-Nicotinamide-adenine (β-NADH), and Adenosine 5’-diphosphoric acid (5’-ADP). On the other hand, several other compounds did not show a significant effect on the growth of *V. mimicus*. In fact, only two compounds, namely 2-deoxyguanosine and 5-hydroxy-L-tryptophan, were found to consistently promote the growth of *V. mimicus* with increasing concentration (Fig. 4 B).

**Fig. 4.**
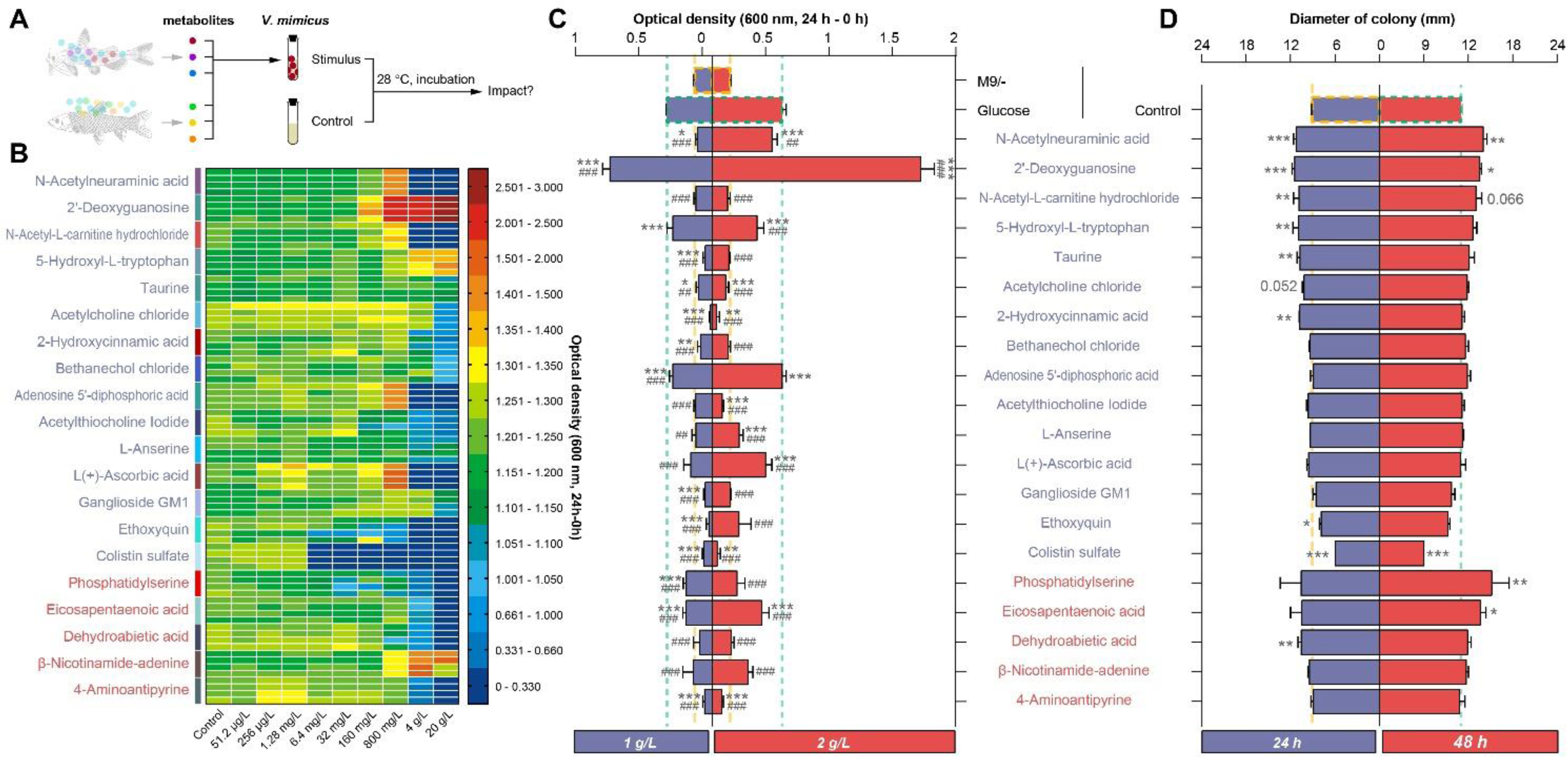
The effects of different compounds on *V. mimicus* life. A: The samples trail for life evaluation. B: the effect of different concentrations of compounds on the growth of *V. mimicus* cultured in LB broth for 24h. C: the effects of compounds as carbon sources on the growth of *V. mimicus.* D: the effect of compound addition on the motility of *V. mimicus*. The significance levels of differences between the M9/– and the M9/compounds group are denoted by asterisks, with * indicating *P* < 0.05, ** indicating *P* < 0.01, and *** indicating *P* < 0.001. The significance levels of differences between the M9/glucose and the M9/compounds group are denoted by hashtags, with # indicating *P* < 0.05, ## indicating *P* < 0.01, and ### indicating *P* < 0.001.

To further expand our investigation, we explored the potential of different compounds to serve as carbon sources for *V. mimicus*. Surprisingly, some metabolites, including 2-deoxyguanosine, N-Acetylneuraminic acid, and 5’-ADP, exhibited efficient utilization as carbon sources, even surpassing the effectiveness of glucose. However, other compounds, like Acetyl L-Carnitine, did not serve as carbon sources (Fig. 4 C). Additionally, we observed that certain compounds, such as N-Acetylneuraminic acid and 2-deoxyguanosine, had a significant impact on the motility of *V. mimicus*, while the effects of other compounds were relatively weaker (Fig. 4 D). Overall, our findings indicate that nucleotide categories, including 2-deoxyguanosine, 5’-ADP, and β-NADH, are particularly favorable to *V. mimicus*. Furthermore, the bacterium shows an attraction to amino acid categories, such as 5-hydroxyL-tryptophan and taurine, while the influence of fatty acyl classes on *V. mimicus* appears to be relatively mild.

### 3.4 Metabolite Changes in the Skin of Yellow Catfish after *V. mimicus* infection

Then, we investigated the impact of *V. mimicus* on specific metabolites identified from yellow catfish. To simulate *V. mimicus* infection, we analyzed the expression of metabolites in the skin tissue (Fig. 5 A). Our analysis resulted in the identification of 265 differentially expressed metabolites out of 836 annotated metabolites. Among these, 32 metabolites were up-regulated, and 233 metabolites were down-regulated, as determined by Student’s t-test (P adjust<0.05) (Fig. 5 B).

**Fig. 5.**
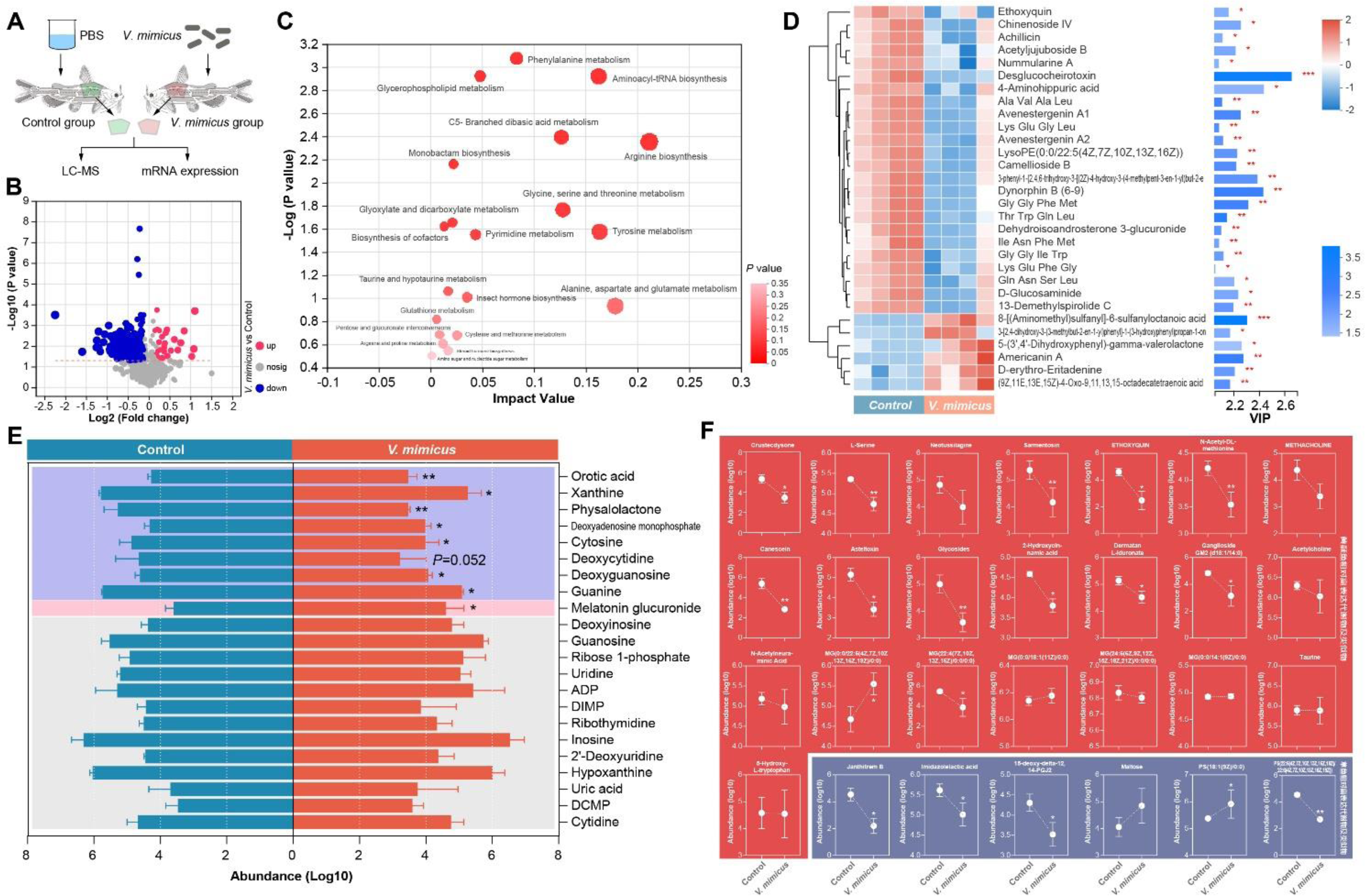
The changes in metabolites in the skin of yellow catfish after infection with *V. mimicus*. A: The samples trail for *V. mimicus* infection. B: the volcano plot of differential expression between groups. C: the top 20 KEGG pathway enrichment results. D: the top 30 VIP analysis of differential metabolites. E: the intergroup expression profile of nucleic acid metabolites. F: the expression of other specific metabolites expressed in yellow catfish and grass carp.

Remarkably, our KEGG enrichment analysis highlighted that the majority of these differentially expressed metabolites were associated with amino acid metabolism, with lipids and nucleotides also exhibiting significant changes in expression (Fig. 5 C). To assess the influence and explanatory power of each metabolite group’s expression pattern, we utilized variable importance in projection (VIP). Oligopeptides were found to constitute a significant proportion of the top 30 metabolites, with 24 of them showing decreased expression in the *V. mimicus* group (Fig. 5 D). Among the 22 identified nucleotide metabolites, eight were significantly down-regulated, while Melatonin glucuronide showed only a slight up-regulation compared to the control group (Fig. 5 E).

Finally, we compared the 28 identified metabolites or analogs between the two fish species. Most of the metabolites that were highly expressed in yellow catfish showed significant down-regulation or a down-regulation trend after infection. Similarly, some metabolites that were highly expressed in Grass carp also exhibited significant down-regulation (Fig. 5 F).

### 3.5 Transcriptional Analysis of Yellow Catfish during *V. mimicus* Infection

To explore the impact of *V. mimicus* infection on the expression of genes related to nucleotide and amino acid metabolism, we conducted a thorough examination of these genes in both skin and muscle tissues. Our investigations revealed intriguing findings. First, *V. mimicus* infection resulted in a significant reduction of 5’-nucleotides in the skin, while a non-significant decrease was observed in the muscle tissue (Fig. 6A). On the other hand, there was a noteworthy decrease in the levels of amino acids in both skin and muscle (Fig. 6B).

**Fig. 6.**
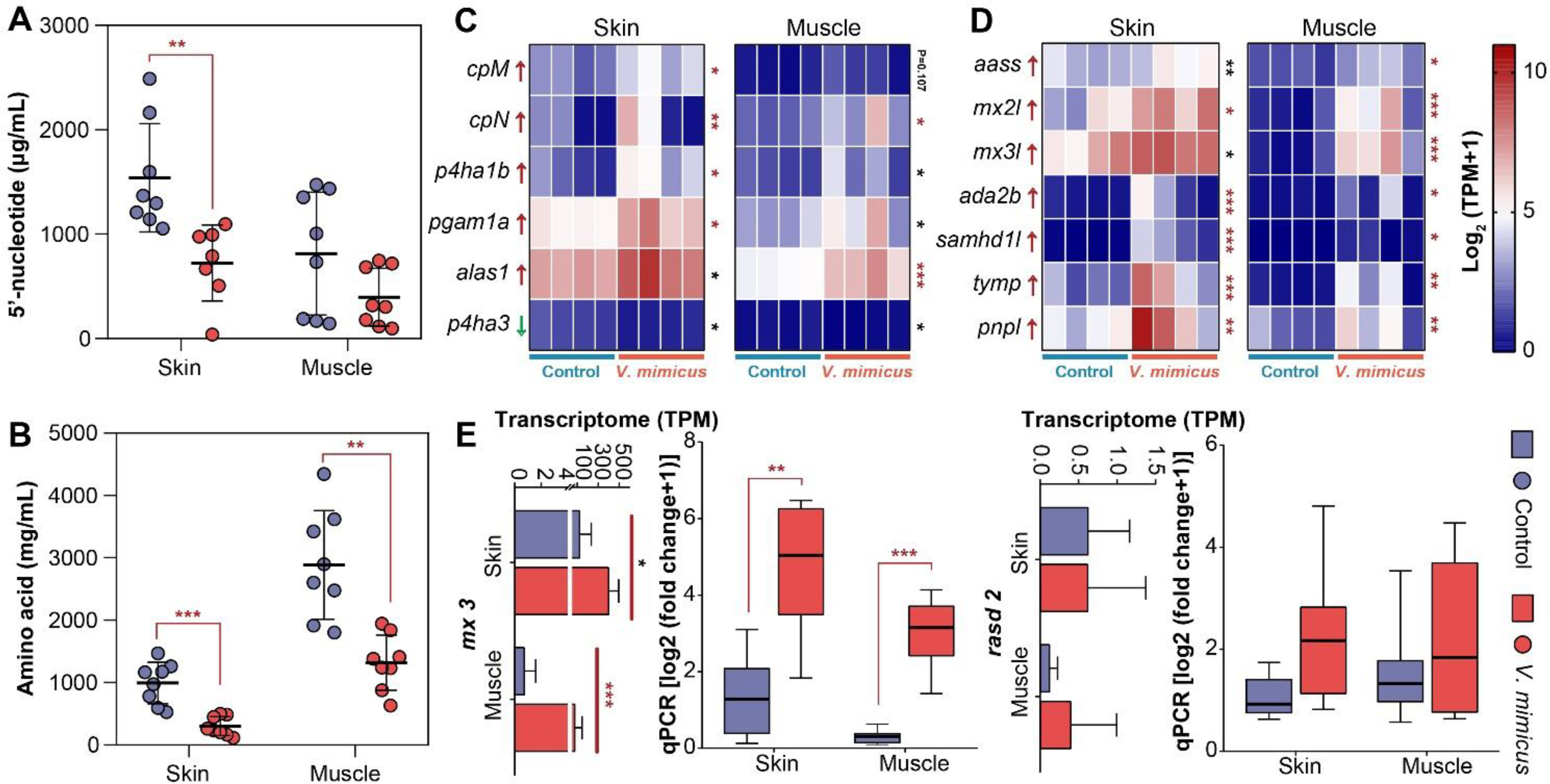
The gene expression analysis of yellow catfish after infection with *V. mimicus*. A and B: the expression of 5’ nucleotide (A) and amino acid (B) between two groups. C and D: the RNA-seq expression of peptidase (C) and nucleotidase (D) related genes between two groups. E and F: the qPCR results of mx-3 (E) and rasd-2 (F) mRNA expression between groups.

Moreover, our transcriptomic analysis yielded valuable insights into the genetic responses to the infection. We observed a remarkable up-regulation of peptidase-related genes, including *cpM*, and a significant down-regulation of *p4ha3*, both playing crucial roles in peptide and amino acid metabolism (Fig. 6C, Table 4). As for nucleotide metabolism-related genes, we discovered that seven nucleotidases, such as *aass*, showed significant up-regulation in both skin and muscle tissues (Fig. 6D, Table 4), and our qPCR analysis further supported the transcriptomic results (Feng et al., 2023)

**Table 4.**
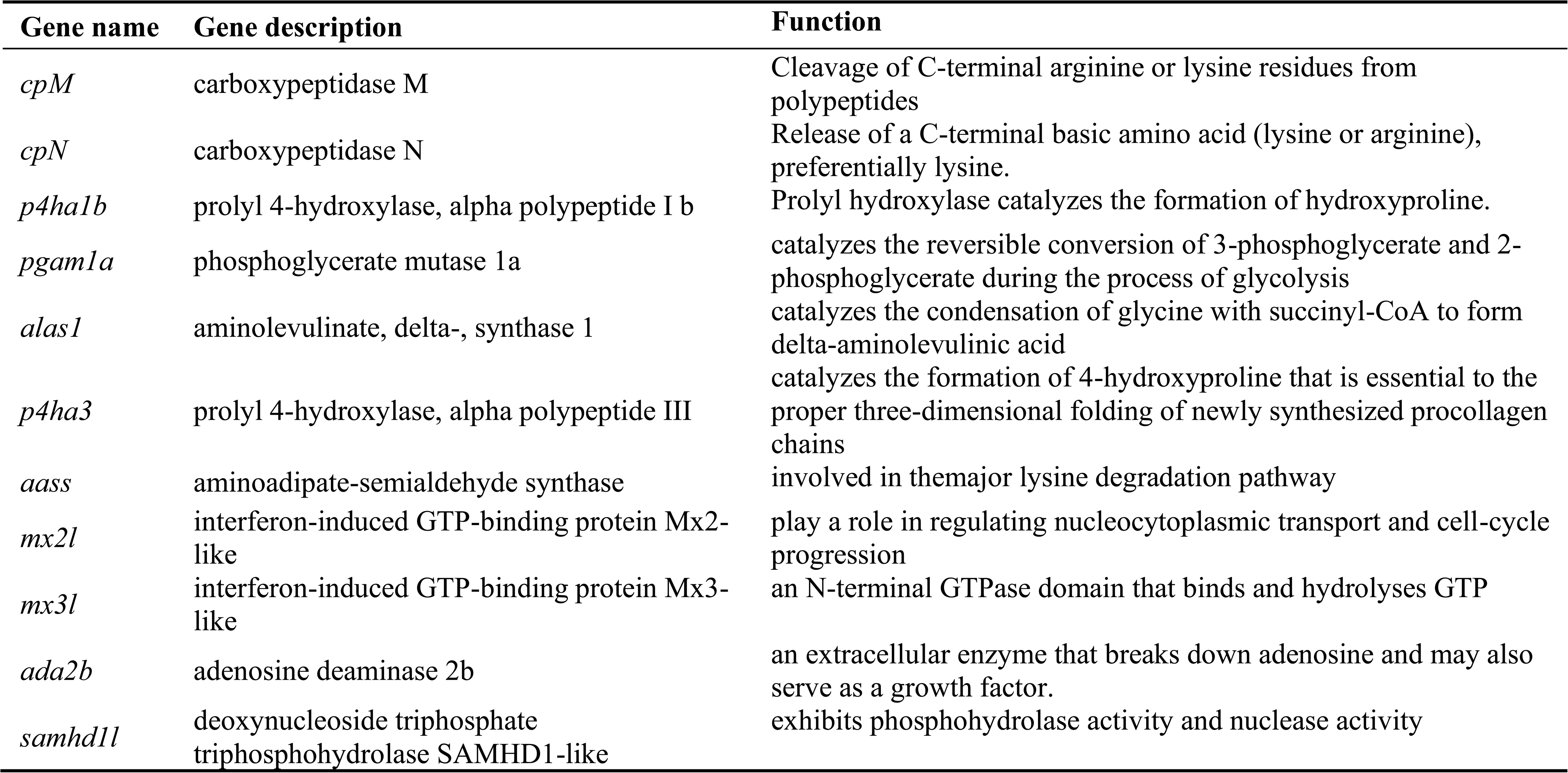
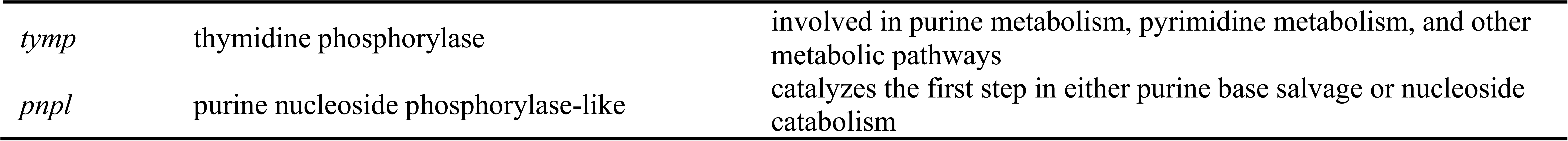
Information on nucleotidase and peptidase-related genes.

Furthermore, we found that *mx-3*, an interferon-induced immune-related GTP-binding protein and GTPase, was significantly upregulated in both skin and muscle tissues following *V. mimicus* infection (*P*<0.05) (Fig. 6E). This finding reinforced the relevance of our transcriptomic data. Conversely, the expression of *rasd2*, another GTP-binding protein with a non-immune function, remained unchanged during infection (Fig. 6F). Lastly, the correlation results were consistent with our transcriptomic findings, adding another layer of confirmation to the observed genetic changes in response to *V. mimicus* infection.

## 4 Discussion

Excluding the immune system, the host functions as a container for bacterial cultivation, and the different nutrient compositions create distinct microenvironments for bacterial survival. Among Siluriformes, yellow catfish are highly susceptible to *V. mimicus* infection, suggesting their unique metabolism facilitates this process. In this study, we selected yellow catfish, the primary cultured fish in China, and the main *V. mimicus*-infected fish in Siluriformes, to investigate the causes of *V. mimicus* super-outbreaks by comparing it with another natural host, Grass carp. Both catfish and grass carp are teleost fish with relatively well-developed immune systems (Smith et al., 2019), similar immune functions (Sahoo et al., 2021), and even immune genes (Chen et al., 2010; Liu et al., 2010; Xiao et al., 2010). As the clinical and pathological symptoms of *V. mimicus* infection are identical between the two fish, this study compared the differences in metabolic composition and potential effects on *V. mimicus* without considering the immune system. Our findings reveal a unique microenvironment dominated by high purine nucleotides and oligopeptides in the skin and muscle of yellow catfish, some of which significantly promoted the growth of *V. mimicus* and were utilized as a nutrient source. During *V. mimicus* infection of yellow catfish, many metabolites were down-regulated, mainly oligopeptides in its skin, and nucleotides and other screened specific metabolites were also significantly weakened. Furthermore, various nucleotidase and peptidase genes were up-regulated in the skin and muscle of infected fish, and the up-regulation of these genes might be related to inflammation. Based on these results, we hypothesize that the high infectivity of *V. mimicus* to yellow catfish may be due to the assisting effect of its specific high purine nucleotide and oligopeptide microenvironment on the body surface, which enhances the growth of *V. mimicus* and thus indirectly enhances the infectivity to the fish. In addition, we previously found that the knocked-out T2SS showed a favorable attenuated effect on the strain, mainly affecting the self-aggregation and metabolism of *V. mimicus* (Yu et al., 2019). *V. mimicus* also showed temperature dependence, and the metabolism genes of *V. mimicus* were significantly inhibited at low temperatures (Kang, 2022). These findings indicate that metabolism plays an essential role in the infection process of *V. mimicus*, causing a significant consumption of metabolites during the infection process.

Based on the results of this study, yellow catfish and grass carp exhibit similar metabolic compositions, indicating that these metabolites could serve as a “baseline medium” for the survival of *V. mimicus*. Certain metabolites may act as “enhancers” to promote the growth of *V. mimicus*, such as nucleotides and oligopeptides which play crucial roles in carbon and nitrogen cycling within organisms. Nucleotides serve as monomeric units for nucleic acid polymers and provide chemical energy, such as ATP and GTP, for numerous cellular functions (Alberts B, 2022). The results of this study also demonstrate that different purine nucleotides exhibit varying degrees of effectiveness in *V. mimicus*. Of the three nucleotide substances selected in this study, 2’-deoxyguanosine was found to be much more effective than the other two. While GTP and ATP are commonly used energy sources in organisms, GMP and AMP serve as basic DNA or RNA structural components. The 2’-deoxyguanosine can be synthesized into dGMP and then converted into GTP, among other molecules (Xi et al., 2000), whereas adenosine diphosphate can be directly converted into ATP and AMP (Xi et al., 2000), which appears to be more efficient than 2’-deoxyguanosine. However, one hypothesis is that different bacteria may exhibit varying degrees of uptake for exogenous nucleotides. For instance, the incorporation rate of 2’-deoxyguanosine is approximately 2.5 times that of thymidine nucleoside, whereas the incorporation rates of deoxyadenosine and deoxycytidine are 0.9 and 0.2 times that of thymidine nucleoside, respectively (Tsuchiya et al., 2020). This could potentially explain why 2’-deoxyguanosine is more effective in promoting the growth of *V. mimicus* than other nucleotides. However, there is currently limited research on the specific mechanisms underlying the effectiveness of 2’-deoxyguanosine, and further studies are warranted in this area.

In addition, peptides, which are short chains of amino acids linked by peptide bonds, are transported through an oligopeptide transport system by most gram-negative bacteria (present in *V. mimicus* SCCF01 genome (Yu et al., 2020)) (Garault et al., 2002). Liu et al. found that deletion of the oligopeptide permease (which facilitates peptide uptake (Kuan et al., 1995)) in *Vibrio alginolyticus* led to a decrease in adhesion, endocytosis, biofilm production, hemolytic activity, and pathogenicity towards *Epinephelus coioides* (Liu et al., 2017). Similarly, the presence of oligopeptide permeases has been found in *Vibrio furnissii* (Wu et al., 2007), *Vibrio harveyi* (He et al., 2011), and *Vibrio fluvialis* (Lee et al., 2004), and has significant effects on the physiological and biochemical characteristics or virulence of these bacteria. It appears that oligopeptides play an important role in the growth or infection process of *Vibrios*, and the high amino acid and oligopeptide environment in yellow catfish may become a preferred nutritional source for pathogenic *Vibrios*. Additionally, in some species, short peptides are absorbed more efficiently than amino acids (Juillard et al., 1995; Liu & Liu, 2020). The uptake of short peptides can also lead to faster synthesis of peptide chains compared to amino acids. This microenvironment of high amino acids and short peptides may increase the efficiency of survival for pathogenic Vibrios and could play an important role in their further entry and pathogenicity in yellow catfish.

In *V. mimicus*-infected yellow catfish, we observed an upregulation of inflammation-induced nucleotidase genes, such as *mx-2*, *mx-3*, *samhd-1*, and *ada2b* (Goldstone et al., 2011; Lorenz et al., 2009; Meyts & Aksentijevich, 2018), while non-inflammation-induced nucleotidases, such as rasd-2, were not differentially expressed. Additionally, histopathological examination revealed significant inflammation in the skin of yellow catfish, and bioinformatics analysis uncovered the presence of multiple inflammatory factors, such as IL-1 and IL-6, in *V. mimicus* infection (Feng et al., 2023). We propose that the inflammatory response eliminates pathogens and dead cells and induces interferon-induced Mx gene expression, thereby influencing nucleotide metabolism in the microenvironment. This process indirectly increases the concentration of purine nucleotides in the microenvironment and promotes the growth of *V. mimicus* (Haller & Kochs, 2002).

Furthermore, while the primary target organ of *V. mimicus* disease in the yellow catfish host differs from that in the human host, *V. mimicus* primarily parasitizes the intestine and causes gastroenteritis in humans (Janda et al., 2015). Nevertheless, similarities between the two hosts exist: the skin of yellow catfish, which is in direct contact with the environment via a mucus layer, is analogous to the human intestine, which is also connected to the external environment via a mucus layer. Skin and muscles are the primary target organs of *V. mimicus* in yellow catfish (Chen et al., 2017), whereas *V. mimicus* mainly parasitizes the intestine (69%) and skin and soft tissues (9%) in humans (Janda et al., 2015). Additionally, the human intestine has a collagen and fibrin environment comparable to skin muscle, a high purine nucleotide environment, and a high oligopeptide environment resulting from food absorption. Therefore, we hypothesize that similar to yellow catfish, the human intestine may also promote the pathogenesis of *V. mimicus* due to the microenvironment formed by purine-rich nucleotides and oligopeptide substances that facilitate the growth of *V. mimicus*. Moreover, *V. mimicus* primarily causes gastroenteritis, similar to atypical *V. cholerae*, with little impact on other organs. Atypical *V. cholerae* strains parasitize a similar proportion of the intestine (56%) and skin and soft tissues (12%) to *V. mimicus*, respectively. In contrast, typical *V. cholerae* strains O1 and O139 parasitize 54% of the intestine (Janda et al., 2015). Thus, the pathogenic mechanisms of *V. mimicus* for hosts may be akin to those of V. cholerae. Studies of V. cholerae’s pathogenic mechanisms have shown that cholera toxin binds to gangliosides on the host’s body surface. The B subunit ring of the toxin attaches to GM1 gangliosides, and the entire toxin complex is endocytosed by the cell. Reducing a disulfide bridge releases the cholera toxin A1 (CTA1) chain, which binds to a human partner protein called ADP-ribosylation factor 6, indirectly over-activating cytosolic PKA and leading to the secretion of H_2_O, Na^+^, K^+^, and HCO_3_^−^ into the intestinal lumen (Wernick et al., 2010). In our study, the cholera toxin genes were annotated in the genome of *V. mimicus* SCCF01 (Yu et al., 2020). Moreover, the skin of yellow catfish is rich in gangliosides, and its catabolic N-Acetylneuraminic acid has a nutritional effect on *V. mimicus*, although less potent than that of purine nucleotides. Therefore, we hypothesize that similar to *V. cholerae*, the abundance of gangliosides on the body surface of yellow catfish may facilitate the adhesion of *V. mimicus* to the body surface (Fig. 7). Ultimately, *V. mimicus* adheres to the host’s body surface through cholera toxin and proliferates in an environment rich in purine nucleotides and oligopeptides, leading to a weakened immune defense, organism morbidity, and possibly death.

**Fig. 7.**
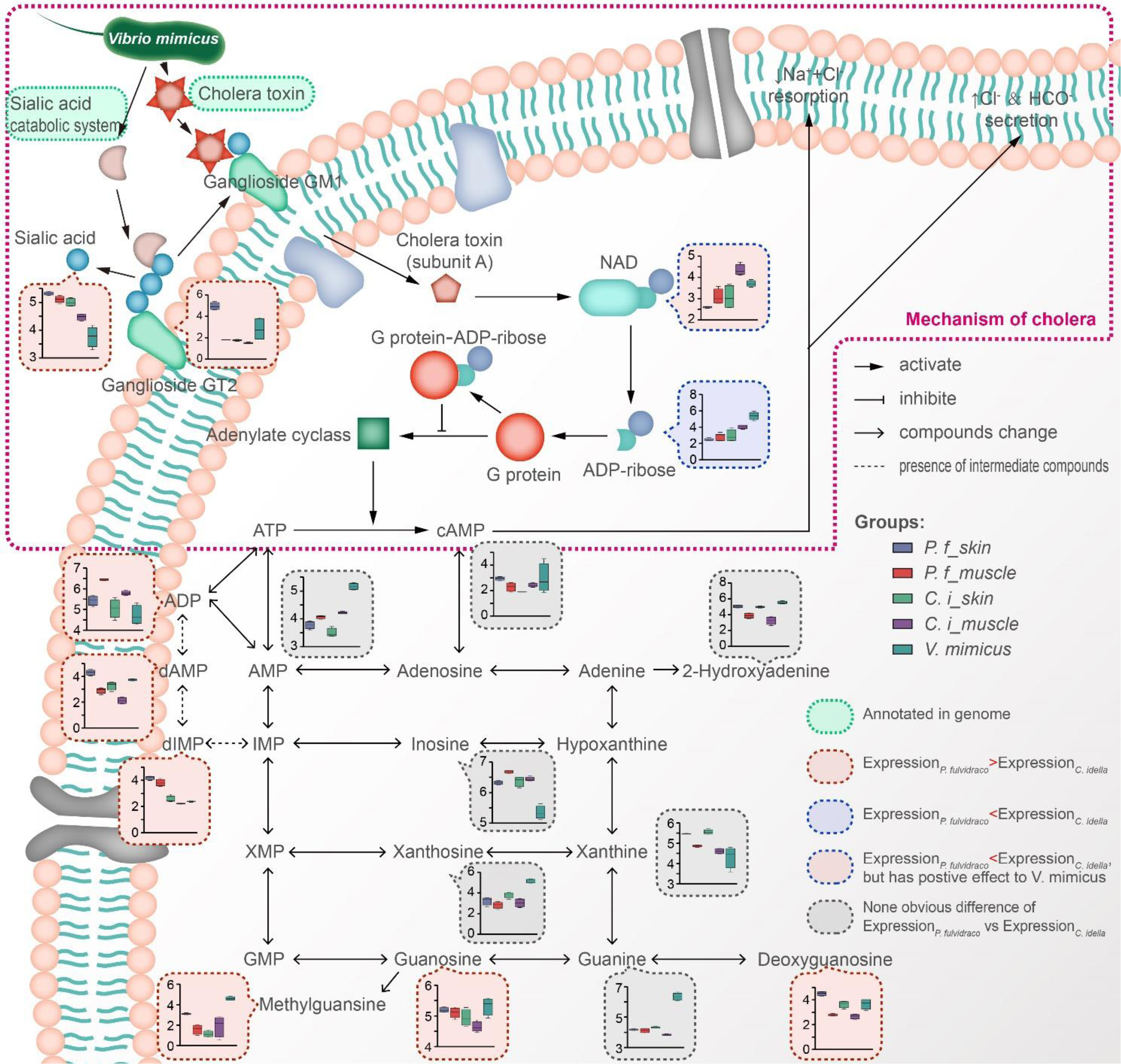
Conjecture on the microenvironment of *V. mimicus* fond of yellow catfish.

## Authors’ Contributions

All authors have read and approved the final manuscript.

## Availability of data and materials

The datasets used to support the conclusions of this study are included in the article.

## Acknowledgments

This study was supported by the Sichuan Province Innovation and Entrepreneurship Technology Talent and Seedling Project (21MZGC0102) and the Sichuan Innovation Team Project of the Agricultural Industry Technology System (No. SCCXTD-15-18). The authors would like to express their gratitude to the funding agency.

## Supplemental Figure Legends

**Fig. S1.**
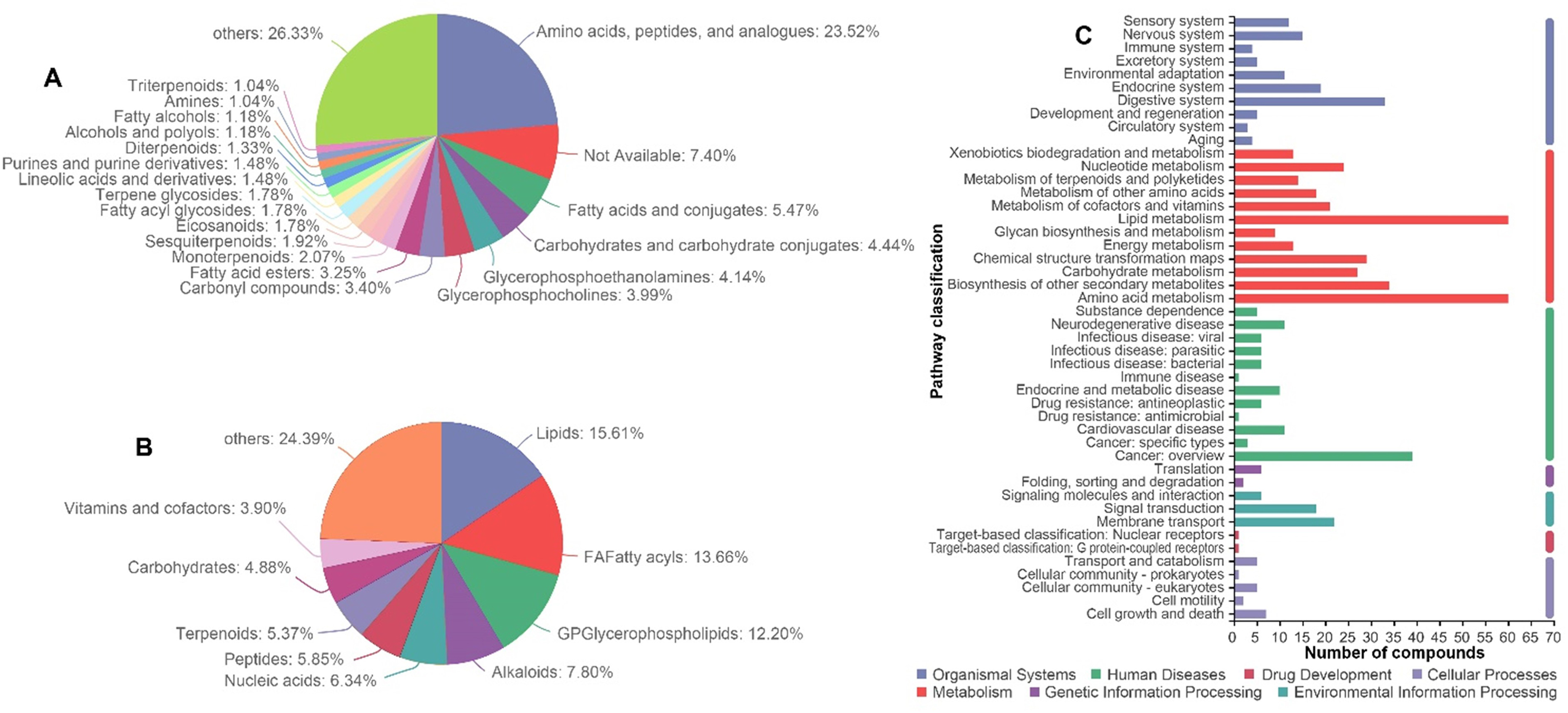
Gross characteristics of metabolite analysis using LC-MS. E: Ions annotated in the HMDB database. F: Ions annotated in the KEGG compounds database. G: Ions annotated in the KEGG pathway database.

**Fig. S2.**
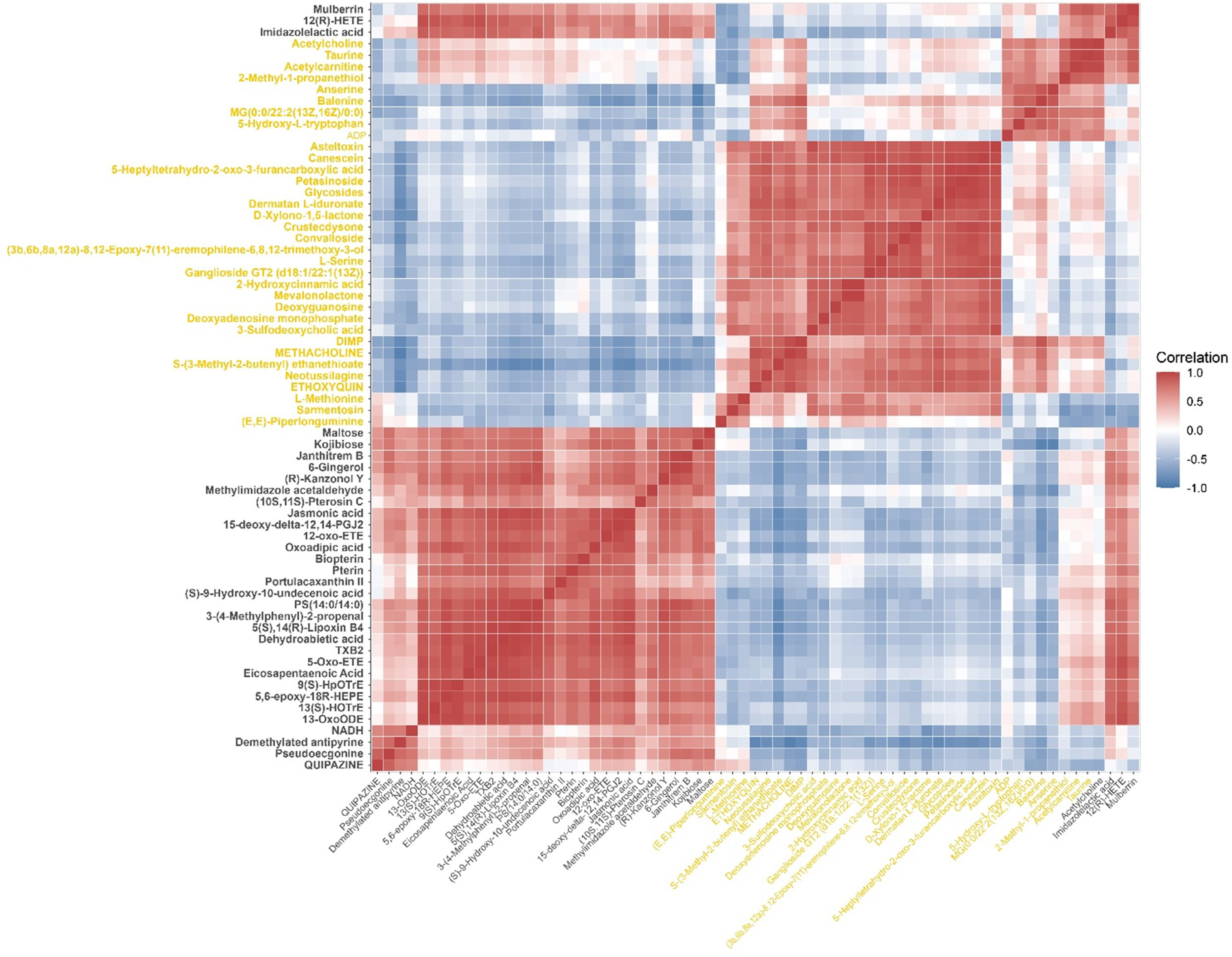
Correlation analysis of 67 target metabolites.

**Fig. S3.**
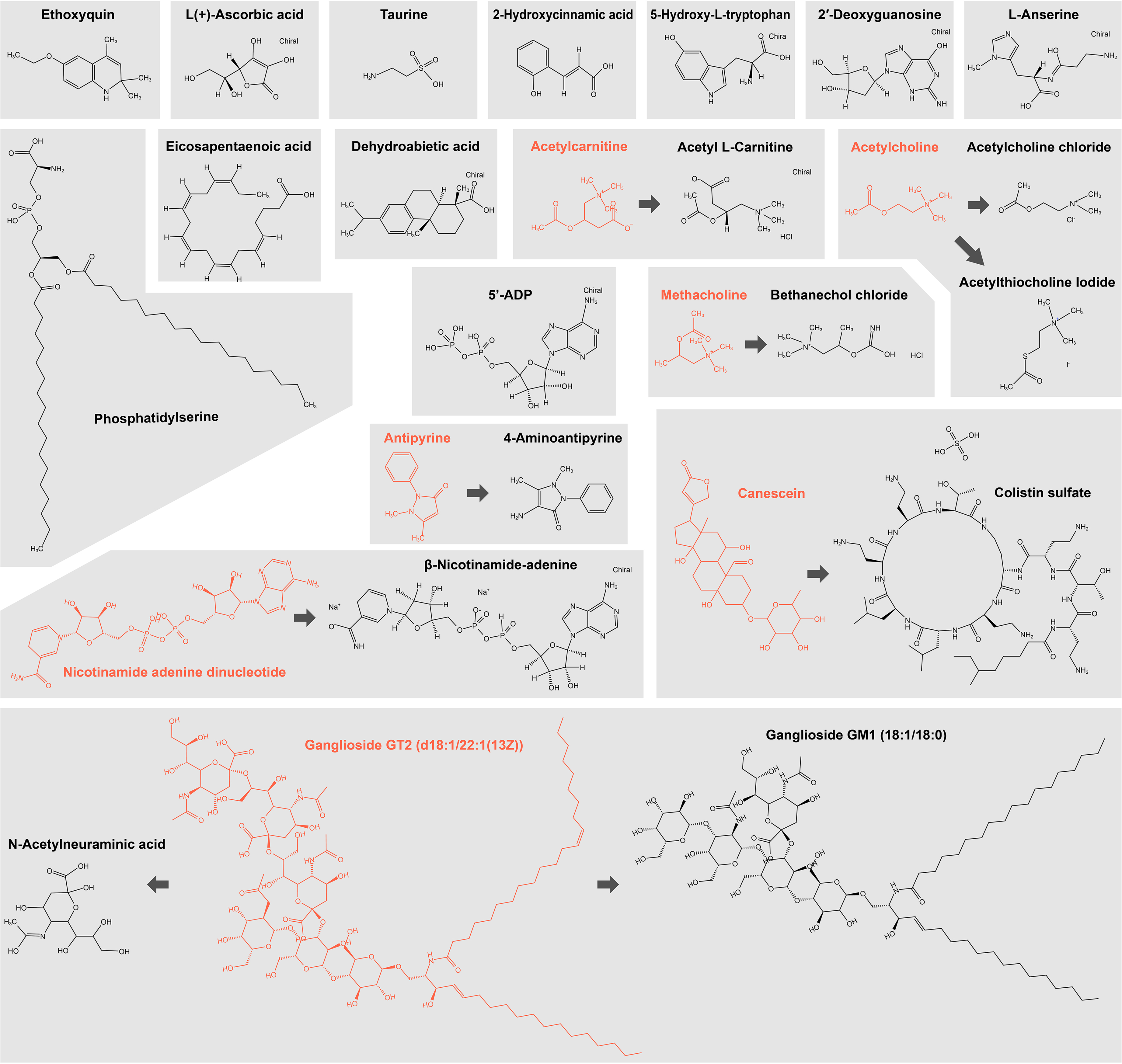
Relationships between candidate compounds and target metabolites and their structural formulas.

